# Decoding activity in Broca’s area predicts the occurrence of auditory hallucinations across subjects

**DOI:** 10.1101/2021.05.21.445102

**Authors:** Thomas Fovet, Pierre Yger, Renaud Lopes, Amicie de Pierrefeu, Edouard Duchesnay, Josselin Houenou, Pierre Thomas, Sébastien Szaffarczyk, Philippe Domenech, Renaud Jardri

## Abstract

**BACKGROUND:** Functional magnetic resonance imaging (fMRI) capture aims at detecting auditory-verbal hallucinations (AVHs) from continuously recorded brain activity. Establishing efficient capture methods with low computational cost that easily generalize between patients remains a key objective in precision psychiatry. To address this issue, we developed a novel automatized fMRI-capture procedure for AVHs in schizophrenia patients.

**METHODS:** We used a previously validated, but labor-intensive, personalized fMRI-capture method to train a linear classifier using machine-learning techniques. We benchmarked the performances of this classifier on 2320 AVH periods *vs*. resting-state periods obtained from schizophrenia patients with frequent symptoms (n=23). We characterized patterns of BOLD activity that were predictive of AVH both within- and between-subjects. Generalizability was assessed with a second independent sample gathering 2000 AVH labels (n=34 schizophrenia patients), while specificity was tested with a nonclinical control sample performing an auditory imagery task (840 labels, n=20).

**RESULTS:** Our between-subject classifier achieved high decoding accuracy (area-under-the-curve, AUC = 0.85) and discriminated AVH from rest and verbal imagery. Optimizing the parameters on the first schizophrenia dataset and testing its performance on the second dataset led to a 0.85 out-of-sample AUC (0.88 for the converse test). We showed that AVH detection critically depends on local BOLD activity patterns within Broca’s area.

**CONCLUSIONS:** Our results demonstrate that it is possible to reliably detect AVH-states from BOLD signals in schizophrenia patients using a multivariate decoder without performing complex regularization procedures. These findings constitute a crucial step toward brain-based treatments for severe drug-resistant hallucinations.

## INTRODUCTION

Hearing distressing voices that other people do not (called auditory-verbal hallucinations, AVHs (1)) becomes a therapeutic impasse for more than 30% of schizophrenia (SCZ) patients (2). These complex sensory experiences are highly variable (3,4), making the characterization of their neurobiological basis especially challenging. This situation creates a technical challenge for people who need to detect/decode AVH states from brain activity for therapeutic purposes.

AVHs were first explored using functional magnetic resonance imaging (fMRI) trait studies, which compared SCZ patients with and without AVHs (5). These studies reported inconsistent alterations in brain connectivity and activity, either increased or decreased, within functional networks associated with language, memory or error-monitoring in patients with AVHs (6,7). One reason for such heterogeneity could reside in the fact that, unlike other symptoms of SCZ, AVHs are intermittent experiences (ON/OFF), an aspect not efficiently addressed with trait designs.

Hence, alternative approaches (called symptom-capture fMRI designs) were developed in an attempt to reduce the inherent complexity of AVHs by focusing on the transient neural changes associated with AVH onsets and offsets (8–11). In these capture studies, hallucinators typically signal AVH occurrence online by pressing a button, revealing a wide range of overactive sensory cortical regions that reflects the high phenomenological interindividual variability in hallucinatory experiences (12–14). Among the brain regions most frequently reported as being associated with AVHs, we can mention the Broca area, the superior temporal gyrus, the temporoparietal junction and the hippocampal complex (11), which are all part of an associative speech-related network more loosely linked with the sensory content of the AVH experience.

However, it remains unclear whether the brain networks identified using online self-report capture designs are (i) involved in the AVH experience itself, or (ii) in the metacognitive/motor processes required to detect and signal the onset of AVHs. Furthermore, these studies referred to a massively univariate activation-based statistical framework known to potentially lose sensitivity by not considering the covariance between voxels (15), which is not particularly well-suited for analyses at the subject level.

In prior studies, we attempted to address these concerns by building upon these paradigms and proposing a button-press free fMRI-capture design based on an independent component analysis, a data-driven multivariate technique (13,16). This approach was proven to reliably detect AVHs from a post-fMRI interview without creating a dual-task situation (i.e., experiencing a vivid AVH and at the same time, pressing a button). However, despite its demonstrated effectiveness, the application of this symptom-capture approach to new patients still required intensive manual labor to individually tailor the analysis pipeline, limiting its translational potential and its applicability/replicability to nonexpert centers. Tackling this issue critically depends on the development of an accurate and easily generalizable method to automatically detect AVHs from brain activity of new SCZ patients with a low additional processing cost.

Here, we propose a novel approach, abstracting out the highly-variable content of AVHs by considering these experiences as stereotyped discontinuous mental events, characterized by core modality-independent properties (17,18). To do so, we combined our previously validated fMRI capture method with supervised machine learning to characterize the informational mapping of AVH states in highly symptomatic schizophrenia patients, both within and between subjects (19). We found that AVH occurrences can be accurately and reliably decoded from individual fMRI BOLD signals and that this signature was robust to concurrent mental processes while being generalizable to new data. This predictive signature is time-selective and appears to mainly rely on the BOLD pattern in Broca’s area. In contrast to previous brain imaging studies that emphasized the distributed nature of AVH brain representations, our results show that a multivariate pattern of neural response in a single hub, namely the Broca area, is sufficient to robustly predict whether a patient is experiencing AVH, paving the way for the therapeutic use of fMRI-based neurofeedback or brain-computer interfaces for closed-loop neuromodulation.

## METHODS

### Participants

Two independent groups of right-handed patients with SCZ (DSM-IV-TR) were recruited and scanned on two different MRI scanners. Twenty-three patients were enrolled in sample 1 (**SCZ#1**) and 34 in sample 2 (**SCZ#2**). They all suffered from very frequent and drug-resistant AVHs (i.e., more than 10 episodes/hour). AVH severity was assessed with the P3-item of the *Positive and Negative Syndrome Scale* (PANSS, (20)). Exclusion criteria were the presence of an Axis-II diagnosis, secondary Axis-I diagnosis, neurological or sensory disorder, and a history of drug abuse (based on a clinical interview and urine tests administered at admission; this did not include tobacco and cannabis use, which were too prevalent in the population). The clinical characteristics of these samples, including the average dosage of antipsychotic medication (in chlorpromazine equivalent, CPZ-Eq), are summarized in **Table 1**. All the patients enrolled in our study provided written consent upon inclusion.

**Table 1.** Demographics and descriptives of the recruited participants.

| Sample (n) | SCZ#1 (23) | SCZ#2 (34) | CTL (20) |
| --- | --- | --- | --- |
| Age (y), mean ± SD | 34.3 ± 8.3 | 35.2 ± 9.8 | 29.9 ± 10.1 |
| Gender, n Female (%) | 11 f (43.5%) | 10 f (29.4%) | 6 f (40%) |
| Duration of SCZ (y), mean ± SD | 16.8 ± 10.5 | 17.1 ± 10.8 | - |
| PANSS-tot, mean ± SD | 79.1 ± 23.3 | 82.4 ± 20.3 | - |
| PANSS-P, mean ± SD | 21.4 ± 5.8 | 21.6 ± 5.5 | - |
| P3-Item, mean ± SD | 5.1 ± 0.9 | 5.6 ± 1.2 | - |
| CPZeq (mg), mean ± SD | 353.6 ± 273.6 | 324.5 ± 246.4 | - |

SCZ: schizophrenia patients ; CTL: control subjects ; y: years ; SD: Standard Deviation ; PANSS-Tot: total score of the Positive And Negative Syndrome Scale ; PANSS-P: positive subscale of the Positive And Negative Syndrome Scale ; P3-Item: 3rd item of the positive subscale of the Positive And Negative Syndrome Scale ; CPZeq: Medication dosage in equivalent - Chlorpromazine (in milligrams).

### Procedure

Each patient underwent a single MRI session after clinical evaluation. This acquisition included two runs: (1) an anatomical MRI and (2) an AVH-capture fMRI. The patients wore headphones and earplugs to attenuate scanner noise and were asked to lie in the scanner in a state of wakeful rest with their eyes closed. The patients reported hallucinatory episodes immediately after the capture fMRI run using a validated post-fMRI interview (13,16,21). This interview allowed us to specify the global frequency of AVHs during the scanning procedure, the relative moment(s) of AVH occurrence and the subjective duration of AVH bouts. These post-fMRI questionnaires were then used to manually identify the ICA components capturing the AVHs for each patient (see the *Data labeling* subsection). If no AVHs occurred during the run, a new capture fMRI session was scheduled. Only patients declaring at least one AVH experience while being scanned were included for further analyses. We followed validated procedures in terms of image parameters and MRI data preprocessing (see ***Suppl.Mat***).

### Data labeling

For each patient, a cortex-based independent component analysis (cb-ICA) was performed. ICA allowed us to blindly extract *N*^th^ components (with *N* equal to 10% of the total number of volumes) from the fMRI BOLD signal time series recorded from cortical voxels. To identify AVH periods, we applied the two-step method summarized in **Fig.1** (13,16,21). Because ICA does not naturally order the resulting components according to their relevance, our first step was to manually sort the ICs capturing only noise or recording artifacts and those capturing neurophysiological sources (IC-fingerprint method). Then, among the components capturing neurophysiological sources, we searched for those with a temporal dynamic compatible with the post-fMRI interview data in terms of number of episodes and times of occurrence. We finally checked whether these ICs contained brain regions previously identified during AVHs (e.g., speech-related network) (see **Fig.1a-b**). As a second step, three labels were defined on the normalized signal time course of these AVH-related ICs (**Fig.1c**): “ON” for the AVH experience (per-AVH periods feature an increased BOLD signal (Z-score > 0) maintained for at least 12 seconds; **SCZ#1:** 2320 **and SCZ#2:** 2000 volumes labeled ON), “OFF” for periods without AVHs (periods with decreased BOLD signals (Z-score < 0) that occurred prior to the “ON” periods and persisted for at least 6 seconds; **SCZ#1:** 997 **and SCZ#2:** 1302 volumes labeled OFF), and “REST” for wider (noisier) resting-state periods, distant from any hallucinatory event (**SCZ#1:** 2974 **and SCZ#2:** 13688 volumes labeled REST, see **Suppl.Fig.1**).

**Figure 1.**
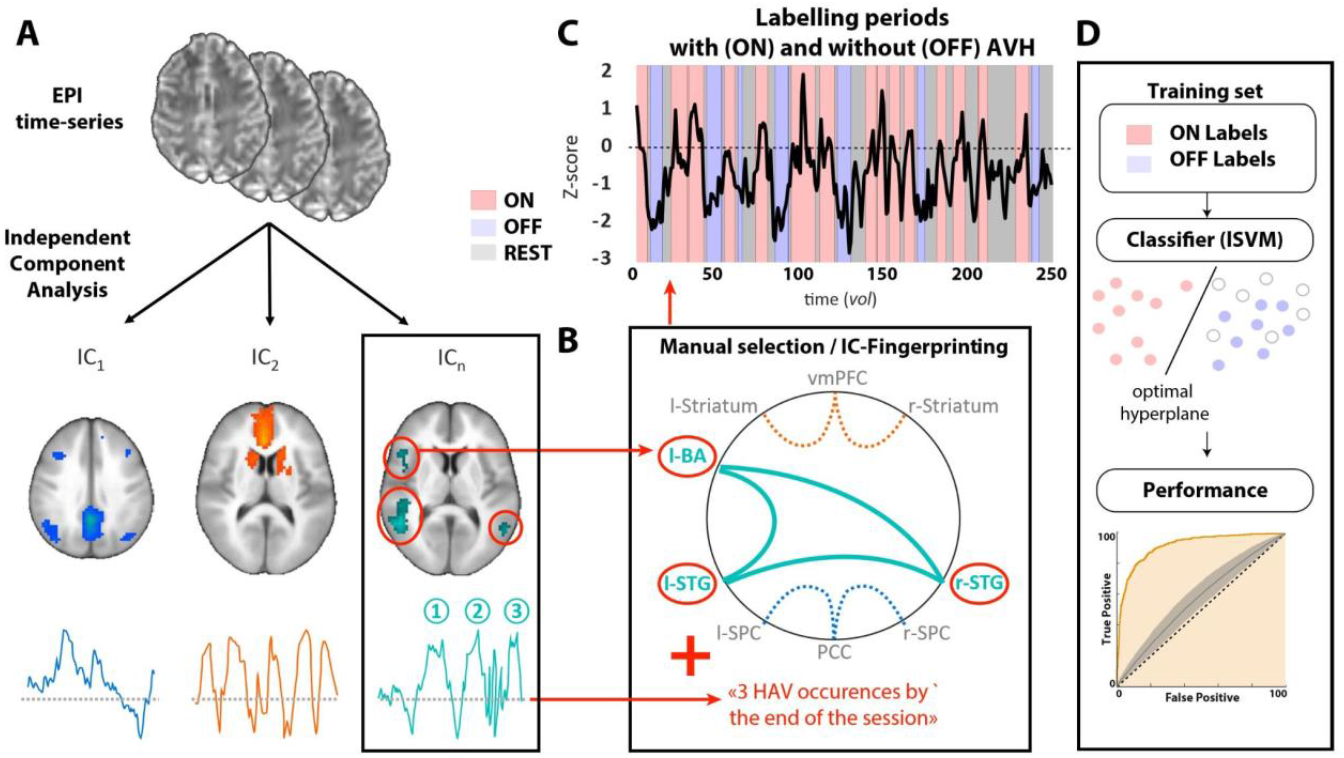
Labeling of the fMRI volumes in ON/OFF/REST periods to train linear classifiers. As previously published (13,16,21), we performed **(A)** a spatial independent component analysis (ICA) of the fMRI time series collected from patients with frequent auditory-verbal hallucinations (AVHs), resting in the scanner (from **SCZ#1** and **SCZ#2** samples). **(B)** IC fingerprinting. For each patient, we selected the component (IC_1-n_) whose spatial network topography matched known AVH-related functional networks (11) and whose temporal activity matched the reported frequency of AVHs based on post-fMRI interviews. **(C)** Finally, we labeled each fMRI volume as being ON (AVH+), OFF (AVH-) or REST, based on the selected component Z-scored temporal dynamics. We labeled fMRI volumes in which the Z-scored component was positive for at least 12 consecutive seconds as being “ON”. Conversely, we labeled volumes as “OFF” if at least 3 consecutive time points were negative and if they were at least 6 seconds away from an ON volume. Finally, we labeled the remaining volumes as “REST”. **(D)** Illustration of the training protocol. ON and OFF labels were used to train a linear support-vector machine (SVM) classifier and find the optimal hyperplane separating these two classes. The independent test set consisted of all voxels, either ON/OFF and REST. EPI: echo-planar imaging; IC_1-n_: independent component 1-to-n; l: left; r: right; vmPFC: ventro-medial prefrontal cortex; BA: Broca area; STG: superior temporal gyrus; SPC: superior parietal cortex; PCC: posterior cingulate cortex.

### Data analysis

We used the Python scikit-learn library to implement linear support-vector machine (lSVM) classifiers (22) using the previously described labels (**Fig.1d**). The workflow described below was conducted independently for each sample (**SCZ#1** and **SCZ#2**).

#### MVPA preprocessing

All fMRI volumes were filtered using an MNI brain mask computed across all subjects. Within this mask, time series were centered and normed. Then, we further reduced the dimensionality of the dataset using an F-test-based univariate feature selection approach. ANOVA parameters were computed for each training set independently of the test set (for each subject, only half of the data were used for training to avoid double dipping). The number of voxels kept for the analysis was approximately 0.5% of the total number of voxels (i.e., 100 voxels). We tested several versions of this filter by varying the number of voxels included (GridSearchCV from 100 to 10,000) and observed that increasing this value had little or no effect on our final results (see **Suppl Table 1.** for an illustration).

#### Supervised analysis with linear SVM

To avoid overfitting and to limit the complexity of our classifier, we used linear SVMs to perform MVPA analyses. Using the labels extracted from the cb-ICA analysis (see *Data labeling* subsection), we trained classifiers to distinguish between MRI volumes labeled ON *vs.* OFF (or between MRI volumes labeled ON *vs.* ALL [OFF+REST]). These classifiers were trained either within subject (using only a subset of the labels from a given subject at a time for training, and testing on the remaining labels) (**Fig.2a-c**) or across subjects (using a subset of the labels from all subjects pooled together and testing on the remaining pooled labels) (**Fig.2b-d**). Corresponding maps are available on Neurovault #. Areas under the curve (AUC) were calculated for each classifier by randomly taking half of the sample data as the training set, and half of the sample data as the test set. ROC curves were computed on the test-set. To assess the statistical significance of the results, we applied two complementary strategies: (i) in the case of within-subject classifiers, we performed a Monte Carlo cross validation with 1000 random 2-fold splits; (ii) in the case of between-subject classifiers, we performed a “leave subject out” cross-validation on a per-subject basis. Hyperparameter optimization is presented in **Suppl.Fig.2**. Discriminative weight-maps illustrating the spatial patterns that best discriminate between AVH states (ON or OFF) were extracted and projected onto glass brains (voxel clusters including more than 10 connected voxels, in the 8-neighbor sense).

**Figure 2.**
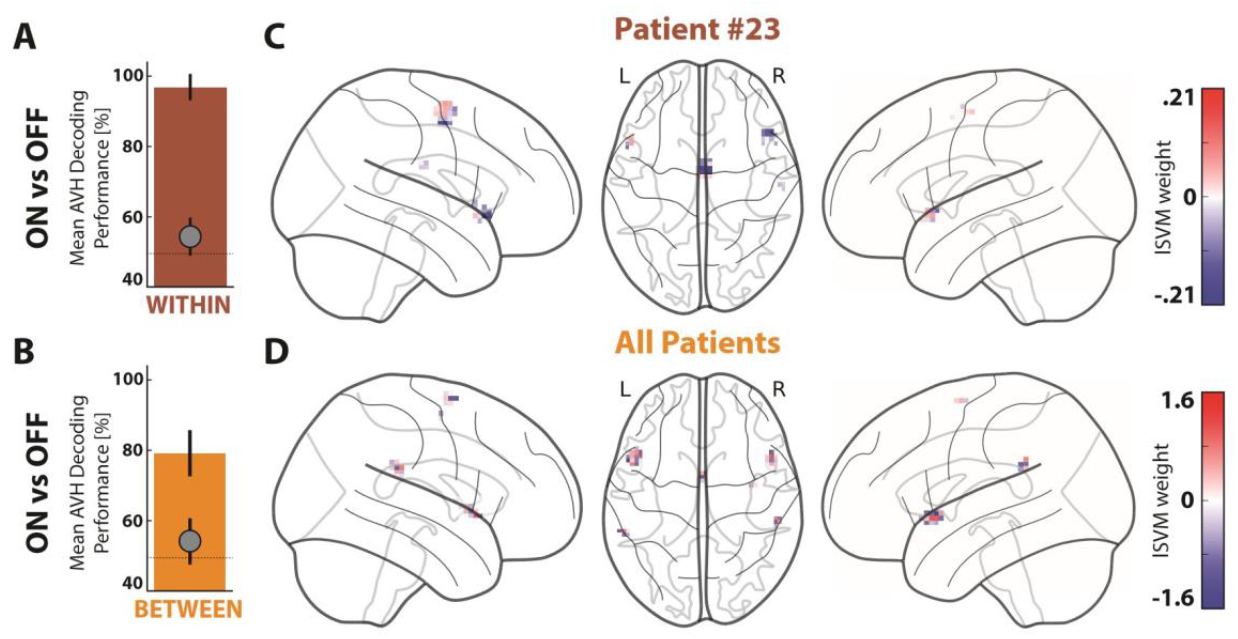
Decoding auditory-verbal hallucinations (AVH) using a linear support-vector machine (lSVM) trained within- (A, C) and between-subjects (B, D). **(A) Group average of within-subject AVH decoding performances** (ON vs. OFF). **(B) Group average of between-subject AVH decoding performance** (ON vs. OFF, **SCZ#1** dataset). The black circle indicates chance level, as estimated using Monte Carlo simulation (with 1000 permutations). Error bars indicate between-subject standard-error to the mean (SEM; panel A-B). **(C) An example of a within-subject contribution map** (patient #23). **(D) Between-subjects contribution map**. The one hundred most informative voxels are color coded to illustrate their contribution to the classifier (lSVM weight). L: left side of the brain; R: right side of the brain.

#### Contribution of local multivariate BOLD patterns to AVH state prediction

To assess the contribution of nonuniform response signs to AVH-state predictions (i.e., the mixture of activation/deactivation in neighboring voxels within a macroscopic region), we ran an additional multiscale sensitivity analysis, in which voxel coordinates were shuffled within the “target” brain regions prior to training and testing lSVMs to decode ON and OFF AVH-states. This procedure intended to destroy all local spatial information within these areas while preserving the target overall voxel BOLD activity distribution. The loss of information induced by shuffling voxel location was quantified by building the corresponding ROC curves and computing AUCs, as was done in the main MVPA analysis.

#### Generalization of the classifiers (I)

The false positive rates of the classifiers were computed using an additional control dataset described below. After a brief training session, 20 healthy volunteers (later called the **CTL group**; mean age = 29.9, min age = 23, max age= 49; see **Table1**) underwent an fMRI experiment on the same 1.5T scanner as the **SCZ#1** sample, during which they performed a “verbal mental imagery” task (8 x 21 s blocks). We chose this condition, as verbal imagery was previously found to share neural correlates with AVHs (23,24). A short verbal instruction (“Imagine”) was given at the beginning of each block, and the participants were asked to stay eyes-closed during the whole acquisition. Data were preprocessed following the same steps as previously described, and functional volumes from this experimental condition were used to challenge the lSVM specificity for AVHs. To take into account the fact that AVH periods typically occur during several time volumes in a row, we convolved the lSVM output probabilities applied to these controls with a flat window of varying size to minimize false positives. The optimal window size was taken as the one minimizing the false positive rates both when the classifier was trained on ON/OFF labels of **SCZ#1** and tested on all volumes of **SCZ#2**, and conversely (see **Suppl.Fig.3**).

#### Generalization of the classifiers (II)

An **out-of-sample cross-validation** step was finally added. Performance generalization of the decoding model built and optimized on the **SCZ#1** dataset was tested using fMRI data from the **SCZ#2** sample kept in a lockbox (25). Conversely, we checked the effect of training/optimizing a decoding model built from **SCZ#2**, later generalized on the **SCZ#1** dataset (kept in the lockbox) for the final performance generalization step.

## RESULTS

We first trained lSVM classifiers to detect BOLD activation patterns concomitant with AVHs, either within subject (one classifier per subject, **Fig 2a-c**) or between subjects (one classifier for the whole **SCZ#1** population, **Fig. 2b-d**). On average, the within-subject classifier used 100 *“ON” vs.* 43 *“OFF”* volumes/subject (see **Suppl.Fig.1**). Its mean decoding accuracy for AVHs was 0.96 ± 0.04 across patients, which was significantly above chance (Monte-Carlo simulations of the null distribution, 1000 permutations; see **Fig.2a**). The between-subjects classifier used a total of 2320 *“ON” vs.* 997 *“OFF”* volumes. Its decoding accuracy for AVHs was also significantly above chance at 0.79 ± 0.06 (1000 permutations, **Fig.2b**). Although MRI data were normalized using only a standard linear transformation, this value was in the same range as the within-subject classifier accuracy. The ROC curve also indicated a high probability for correct classification for the between-subjects classifier (AUC_H0_=0.55 *vs*. **AUC=0.85**, **p<1e^-3^; Fig.3a**).

**Figure 3.**
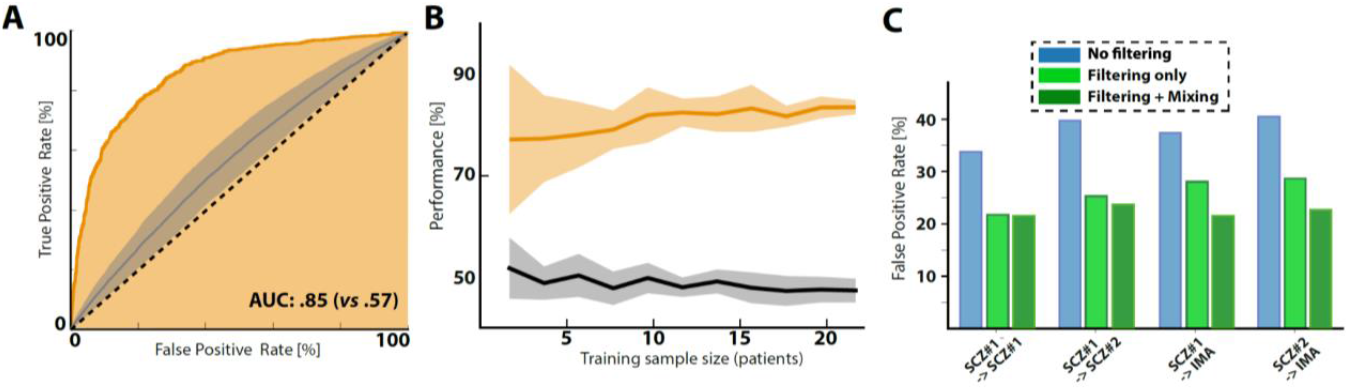
Reliability and performances of between-subject decoding of auditory-verbal hallucinations (AVHs) using a lSVM classifier (ON *vs*. OFF). **(A) The receiver-operating characteristic curve** indicates the trade-off between false positive and true positive label classification (± standard error to the mean, SEM). **(B) Reliability of between-subjects AVH decoding performance.** Mean between-subjects decoding performance (± SEM) as a function of the number of patients in the training set (red line). The black line indicates the chance performance level (± SEM) estimated using Monte Carlo simulation (1000 permutations). See also **Suppl.Fig.4** presenting a correlation analysis between performances and the number of ON periods **(C) False positive rates** of a between-subjects lSVM trained on the **SCZ#1 (or SCZ#2)** dataset (ON vs. OFF labels) and then applied to the **CTL** dataset (could be **SCZ#1 or SCZ#2** with ON vs ALL labels, or **IMA** where IMA is a third dataset collected while performing a verbal mental imagery task (see Methods)). False positive rates are shown for raw data (blue bars) when the lSVM scores are filtered, i.e., convolved with a flat time window of 5 s to ensure that ON periods are contiguous (orange bars; see Methods), or when, in addition to this filtering window, half of the test data are also used to refine the training sets.

Then, we built contribution maps from the predictive weight given to each voxel by the “ON” *vs.* “OFF” lSVM classifier, which revealed AVH-related BOLD activity patterns in the bilateral inferior frontal gyri (i.e., the Broca area and its right homolog, hereafter called BA), the supplementary motor area and the pre-SMA (“SMA” will be used to designate these two structures), and the bilateral supramarginal gyri (**Fig.2c-d**). These brain regions have previously been shown to be involved in AVH pathophysiology (6,11,23).

As a sanity check, we ensured that these performances did not depend on the ON/OFF volume ratio across subjects, or on frame displacements (**Suppl.Fig.4**). We confirmed that these findings and the performance of the classifier did not depend on the threshold applied for feature selection (i.e., the number of voxels included in the training-set up to 10,000 voxels; see **Suppl.Table 1**) and that between-subjects decoding performances were stable and reliable for all training sample sizes larger than 10 patients, indicating that the multivariate pattern of BOLD activity used by the classifier seems especially robust and well conserved between subjects (**Fig.3b**).

We also ran a series of additional analyses aimed at assessing the specificity and robustness of this between-subject ON *vs.* OFF classifier. First, we estimated its false positive rate when applied to new data. We applied the classifier trained with **SCZ#1** ON/OFF labels to noisier volumes (ON *vs.* [OFF + REST]) either taken from the same dataset (**SCZ#1**, n=23), or from a different dataset (**SCZ#2**, n=34). As shown in **Fig.3c (blue bars)**, false positive rates were initially high in both cases (i.e., 35% on average). However, by applying a smoothing kernel to the output of the classifier to take into account the fact that ON and OFF periods should cover consecutive volumes, the false positive rate dropped at 21% (with an AUC of 85% for **SCZ#1** -> **SCZ#1** and 81% for **SCZ#1**->**SCZ#2**). It is noteworthy that the application of the smoothing kernel mainly acted by limiting the risk of FP over the 3 volumes preceding or succeeding actual AVH states (**Suppl.Fig.3c-d**).

Similarly, when applying the classifier to a third dataset acquired in healthy control subjects performing a verbal imagery task (a mental process potentially overlapping AVH; see **CTL** dataset, *Methods* section), the initial false positive rates were high (37%), but filtering the classifier’s output resulted in decreasing this false positive rate to 25% (**Fig.3c, light-green bars**). Finally, retraining the lSVM classifiers on a composite dataset mixing “OFF” labels with some REST and/or IMA labels from the #SCZ1/2 and/or IMA dataset (i.e., using a subpart of the **CTL** dataset to enhance the training sets) reduced false positives to a reasonable level of approx. 20% in all conditions without altering AVH decoding accuracy (**Fig.3c, dark-green bars**; see also **Suppl.Fig.5** presenting the decoding accuracy and contributive maps for a between-subject classifier trained on ON *vs.* [OFF + REST] with **SCZ#1**).

To ensure out-of-sample generalization and control for the risk of hyperparameter overfitting, the classifier trained on the **SCZ#1** dataset was then challenged to predict the labels of the independent **SCZ#2** sample (fully held-out from the optimization process and only accessed to generate a new unbiased estimate of the AVH decoder’s performance). We switched this procedure by training a classifier on **SCZ#2** and predicting labels on **SCZ#1**. In both cases, the AUCs were found to be significantly different from chance (0.85 and 0.88 respectively), while the resulting contributive voxels overlapped in 75% of the cases within the BA (**Fig.4**).

**Figure 4.**
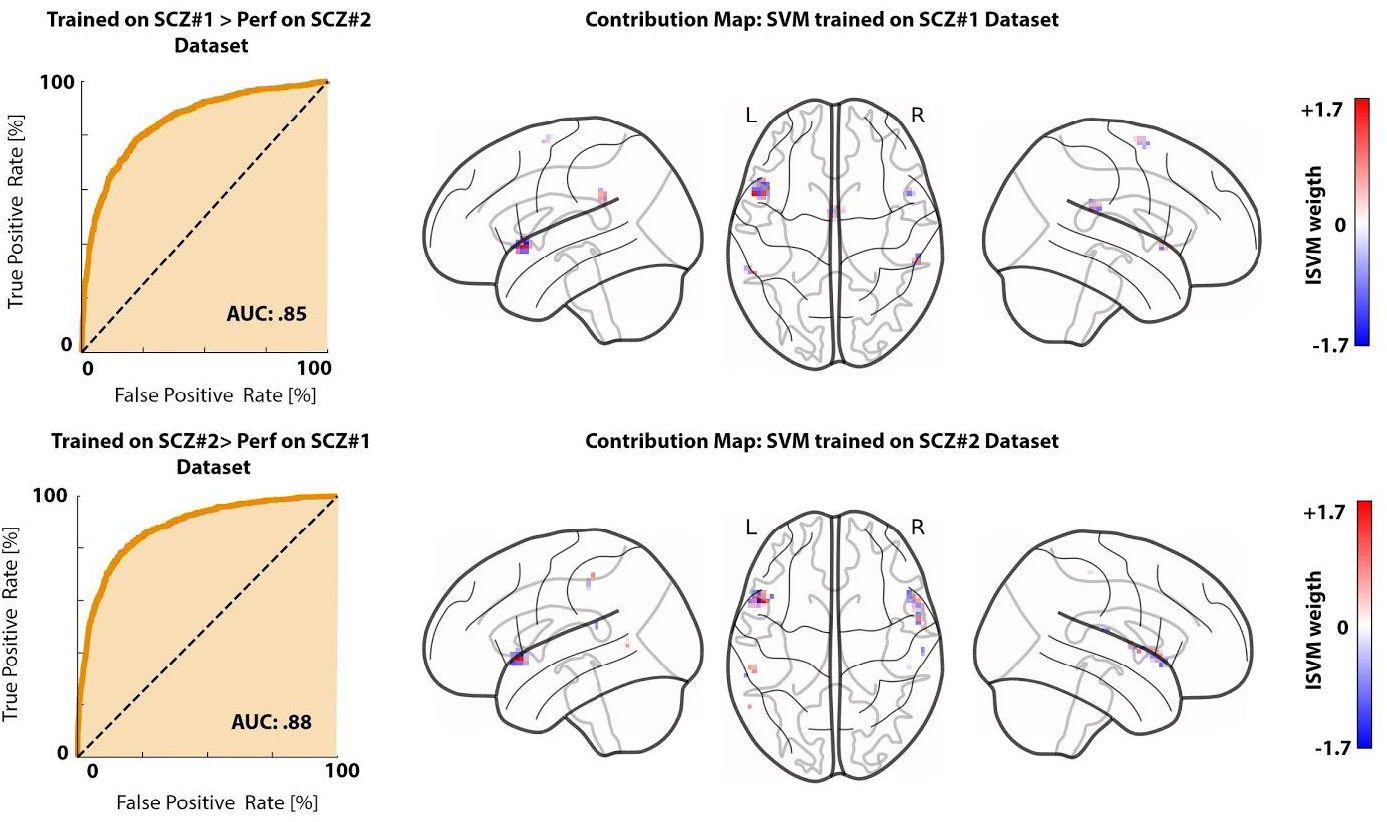
Generalization of the AVH decoder to out-of-sample data. We checked for possible overfitting and overhyping of the AVH linear support-vector machine classifier (lSVM). In conjunction with the cross-validation approach used on the **SCZ#1** dataset (cf. **Figs.2-4**), we refer to a lockbox approach, in which the second sample of SCZ patients scanned during AVH occurrences was set aside from the beginning of the study and used only after hyperparameter optimization of the AVH classifier to check its accuracy on a completely new dataset (upper panel). We repeated the procedure to ensure that we could obtain similar generalization performances when starting training and optimizing the decoder on the **SCZ#2** dataset and testing accuracy on the **SCZ#1** dataset (lower panel). L: left side of the brain; R: right side of the brain.

Finally, we assessed how much AVH-related information was local, encoded as a spatial mixture of activation/deactivation within each brain region identified in the main analysis (namely, the BA and SMA) while controlling for global patterns of average regional levels of BOLD activity (for example, higher mean BA BOLD activity and lower mean SMA activity during AVHs). We found that shuffling voxel coordinates within the BA and SMA prior to training/testing the classifier had a significant effect on AVH prediction within the BA (**Fig.5a**, blue ROC curve, **p < 1e^-5^**), but not within the SMA (**Fig.5a**, pink ROC curve). Moreover, ROC curves when only BA voxels were shuffled were not statistically significant from the ROC curves when both BA and SMA voxels were shuffled (**Fig.5a**, dashed black ROC curve). This suggests that most AVH-related information captured by the linear classifier is represented locally in the spatial BOLD pattern of Broca and not elsewhere in the brain.

**Figure 5.**
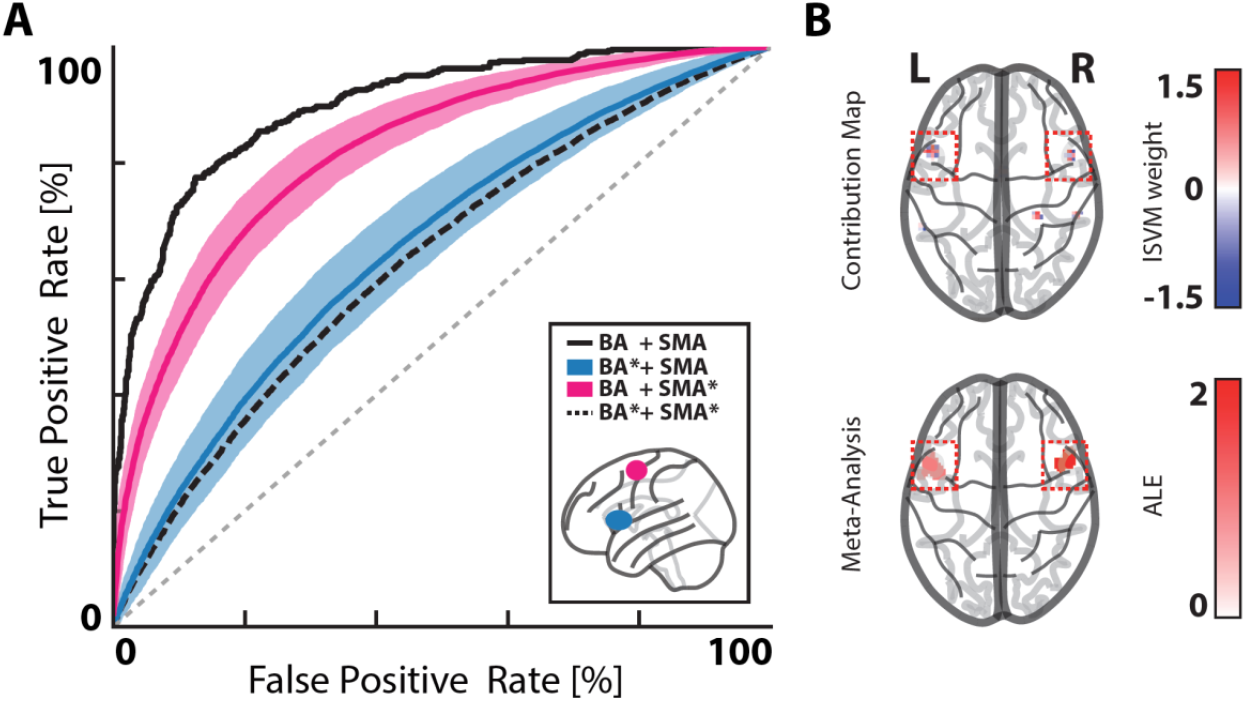
Local *vs* global patterns of BOLD activity associated with AVHs. **(A) Receiver-operating characteristic curve of a between-subjects ON vs. OFF lSVM** indicating the trade-off between false positive and true positive label classification (black line, **SCZ#1** dataset). Pink and blue lines indicate the receiver-operating characteristic curves after a volume-by-volume shuffling of voxel locations within each of the two regions-of-interest (ROIs, pink: supplementary motor area (SMA) shuffled, blue: Broca area (BA) shuffled). The dashed black line indicates the performance of the classifier when the spatial locations of voxels within both ROIs are simultaneously shuffled. Note that voxels belonging to an ROI are always shuffled to a location within that same ROI. Shaded blue and pink areas indicate between-subjects standard-error the mean (SEM). **(B) Contribution map of the between-subjects lSVM** (ON vs. OFF; top panel) **against an updated activation likelihood location (ALE) meta-analysis of the literature on per-AVH BOLD activity** (bottom panel). L: left side of the brain; R: right side of the brain.

Taken together, these results shed new light on the pathophysiology of AVHs, which was previously thought to be inherently distributed in a complex network of brain-wide regions, and suggest instead that a nonuniform BA functional pattern is critical to predict AVH occurrences and set it apart from normal perception.

## DISCUSSION

The online detection of spontaneous AVH occurrence has long been very challenging (9,13,16). By demonstrating that a simple linear classifier can robustly decode hallucinatory states from out-of-sample resting-state fMRI data without complex preprocessing, we substantially extended recent work in the field of AVH fMRI capture by departing from a classic activation-based to a multivariate information-based perspective. Notably, we found a 79% between-subjects accuracy in distinguishing hallucinatory and nonhallucinatory periods (ON *vs.* OFF, .85 AUC).

We demonstrated the robustness of our findings by conducting a set of supplemental analyses to precisely characterize our decoder features: (i) repeating the performance measures while increasing the sample size, (ii) confirming good performances even when using noisier data, and (iii) reducing false positive rates after enforcing temporal regularization and training the algorithm to selectively ignore verbal imagery. We further addressed a crucial issue in the classification literature: replicability of the decoder performances (26). Here, we went beyond conventional leave-one-subject-out cross-validation strategies by replicating our results both ways, using either SCZ#1 or SCZ#2 as a training/test set or as an out-of-sample dataset aside in a lockbox (and thus independent from the optimization process (25)).

We identified a BOLD multivariate signature predictive of AVH in speech-related motor/planning brain regions, such as the Brodmann area 44, part of the BA (27), and the SMA (Brodmann area 6, medially (28)). This functional signature coincides with previous per-AVH activation reports in first-episode psychosis (13,29), schizophrenia patients (9,16,30,31) or nonclinical voice hearers (23). Interestingly, this last study reported differences in the timing of SMA activations (relative to BA activations) between AVHs and a verbal imagery condition, which appears fully compatible with our optimization procedure that allowed us to strengthen AVH/imagery discrimination. Additionally, consistent with these findings, our BA cluster overlaps with coordinate-based meta-analytic findings of per-AVH hyperactivations in schizophrenia (11) or conditioned hallucinatory mapping obtained from nonclinical participants (32) (**Fig.5b**, see also **Suppl.Tables 2 & 3, Suppl.Fig.6**).

The BA and SMA are also known to be involved in error monitoring and inhibition (33), suggesting that AVHs may result from aberrant motor representations/predictions (despite an absence of online self-report in the participants), which may be a core mechanism in the lack of insight typically associated with hallucinations (34). This appears compatible with prior hypotheses of inner speech as a form of action (35). In contrast, hippocampal or temporoparietal structures, also known to be involved in AVH pathophysiology (6,36–38), possibly by reflecting the spatiotemporal, rich and complex content of these experiences (21,39), were not necessary to reach high decoding performances. Even if anteroposterior dysconnectivity between speech-related areas has been regularly shown to be involved in AVHs (40), this new finding suggests either that (a) the highly variable nature of the information computed by these temporal-hippocampal structures is not stereotyped enough to be decoded using lSVM or (b) that most of the fine-grained relevant information conserved between subjects is encoded in Broca’s BOLD activity.

Although the main goal of this study was to demonstrate the feasibility of a reliable and easily deployable multivariate AVH decoder, it also adds several insights to AVH pathophysiology. We know that the performance of a classifier is dependent on the functional features used to train the lSVM. This is why we referred to a valid strategy to determine ON/OFF labels (13,16,21) that proved able to achieve good performances even without special regularization preprocessing steps (41). This may appear surprising at first glance since previous work conducted on more subtle functional profiles (i.e., states preceding AVH onsets) showed that specific classification algorithms with total variation penalty were better at detection than lSVM (39).

In reality, the good performances demonstrated by our classifier reflect the remarkable consistency of the AVH-related BOLD pattern across patients, robust to varying magnetic field strength or sequence parameters (e.g., image resolution, TR or differences in number of volumes between **SCZ#1** and **SCZ#2** datasets). Limited activation studies have previously reported similar spatial stability in per-AVH activation patterns (16,30), and we propose that such consistency could be due to the involvement of the BA.

Broca’s area is a highly preserved anatomical-functional hub (for an evolutionary perspective, see (42)), which may relate to the coding of a very generic, amodal, feature of the AVH experience. In this vein, the BA could be sensitive to intrusiveness into consciousness, irrespective of the highly variable phenomenological content of the voices (4). Even if speculative at this stage, this assumption appears compatible with recent findings showing the involvement of the inferior frontal gyrus in the intrusion of unwanted thoughts more broadly, notably in OCD patients suffering from severe obsessions (43), while states of “mind-blanking” were shown to be associated with BA deactivation (44).

In our study, this assumption was also confirmed by voxel permutation tests performed to challenge the local distribution of response signs in contributive maps. Such permutations flattened the lSVM accuracy when applied to the BA, while extending this operation to the SMA only slightly (yet significantly) impacted the classifier. This can be interpreted as a form of redundancy in the information processed by the SMA, while the BA could locally compute crucial elements for AVH intrusion prediction, coded in its microstructure (and not elsewhere in the brain), only experimentally accessible due to multivariate pattern classification methods.

Until now, AVH fMRI capture has been considered to be complex and time-consuming because of its many technical constraints. This situation has limited the use of these methods to offline applications in the lab, which has significantly hindered therapeutic innovations. Thanks to our newly validated and replicable biomarker, reading out hallucinatory states online from resting-state fMRI is now possible, providing a gateway to further validate fMRI-based neurofeedback procedures to relieve severe AVHs and develop brain-computer interfaces for closed-loop neuromodulation (45). Indeed, both approaches require clearly defined cortical targets at key points in the brain networks involved in AVHs (46).

The unpredictable nature of brain-state changes over time associated with hallucinations has long remained a major (and supposedly insuperable) challenge in neuroscience, and most therapeutic alternatives to medications have attempted to modulate network activity (see for instance, noninvasive brain stimulation targeting the temporoparietal junction (47)). We believe that our findings not only uncovered a neurofunctional reconfiguration associated with this fascinating mental experience but also provide a way to automatically identify a dynamic neural pattern playing an important, if not critical, role in AVH occurrences (i.e., intrusiveness), paving the way for the generalization of fMRI capture and the development of new image-guided therapeutic strategies for drug-resistant hallucinations.

## DECLARATIONS

### ETHICS APPROVAL

The study received approval from a national ethical committee (CPP Nord-Ouest France IV, #2009-A00842-55), and written informed consent was obtained for each participant enrolled in the study.

### AUTHOR CONTRIBUTIONS

The project was supervised by RJ. Data was collected and prepared by TF and RJ. Data were analysed by PY, PD and RJ. All the authors contributed to the data interpretation and paper writing.

### AVAILABILITY OF DATA

Resulting maps from this study will be made available of a public depository after acceptance (like NeuroVault): TO ADD

## ACKNOWLEDGMENTS

This study was supported by the *Agence Nationale de la Recherche* (ANR-16-CE37-0015 INTRUDE awarded to RJ).

## SUPPLEMENTARY MATERIAL

### I. Supplementary Methods

#### Imaging parameters

Patients from the **SCZ#1** sample were scanned on a 1.5T MRI scanner, while patients from the **SCZ#2** sample were scanned on a 3T scanner (both Philips Achieva series). The anatomical run was identical for all the participants and consisted of a T1-weighted 3D multishot turbo-echo scan (150 transverse slices, field of view = 256 mm^2^, voxel size = 1 mm^3^).

The capture fMRI run differed between the two groups. For **SCZ#1**, we acquired a set of 280 blood oxygen level-dependent (BOLD) volumes (single-shot sensitivity-encoded EPI sequence, 30 transverse slices, field of view = 240 mm^2^, voxel size = 4 mm^3^, repetition time = 3000 ms, echo time = 70 ms, total acquisition time = 14 min) and 900 BOLD volumes for **SCZ#2** (3D-PRESTO SENSE, 45 slices, field of view = 206 mm^2^, voxel size = 3.3 mm^3^, dynamic scan time = 1000 ms, echo time = 30 ms, total acquisition time = 15 min).

#### MRI data preprocessing

Data-preprocessing was performed with the BrainVoyager software suite (v21, Brain-Innovation, Maastricht, NL). Functional data were submitted to:

i. a slice scan time correction (only for the **SCZ#1** sample, since **SCZ#2** data were acquired in volume)
ii. 3D motion correction for head movements using a rigid-body algorithm (frame displacements were extracted to run supplementary control analyses)
iii. smoothing, using a spatial Gaussian filter (full-width at half-maximum [FWHM] = 6,0 mm)
iv. temporal high-pass filtering with 2 sin/cos (i.e., two sine waves fall within the extent of the data).

Anatomical data were submitted to an intensity inhomogeneity correction, while anatomical-functional coregistration was performed through white-grey matter boundary-based alignment (48). After quality check, all anatomical and functional volumes were spatially normalized to the MNI space.

### II. Complementary description of the datasets/ labels

**Supplementary Figure 1.**
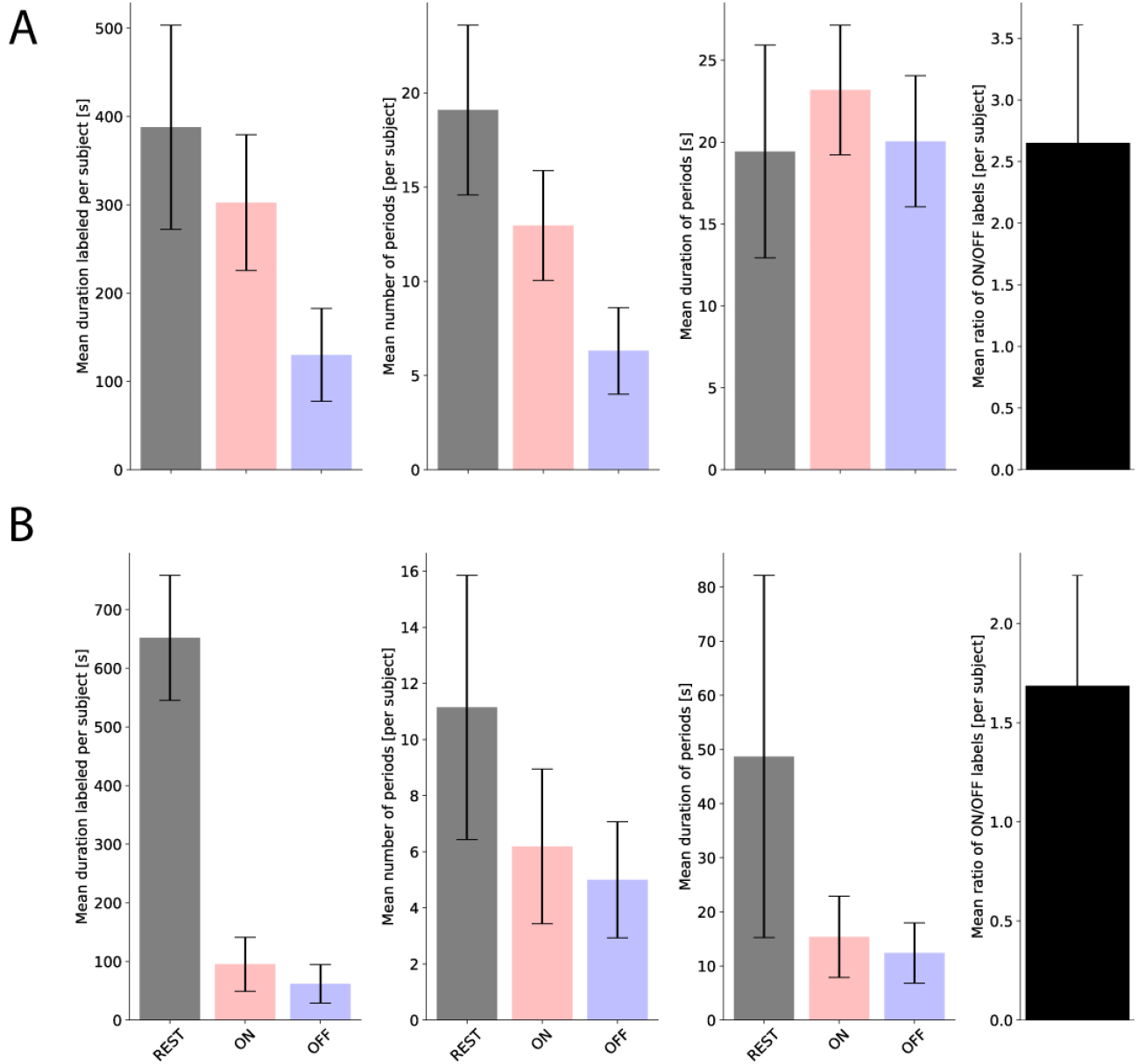
Distribution of the ON/OFF/REST labels for hallucinations in the datasets. **(A)** Mean duration (in seconds) for the ON, OFF and REST periods per subject of the **SCZ#1** sample (left panel), the mean number of periods with contiguous labels per subject (center), and the mean duration of these ON/OFF/REST periods (right panel, in seconds). The last panel on the right shows the average ratio of ON/OFF/REST labels per subject. Error bars show the standard deviation. **(B)** Same as panel (**A**), but for the **SCZ#2** sample.

### III. Sanity-check analyses

**Supplementary Table 1.**
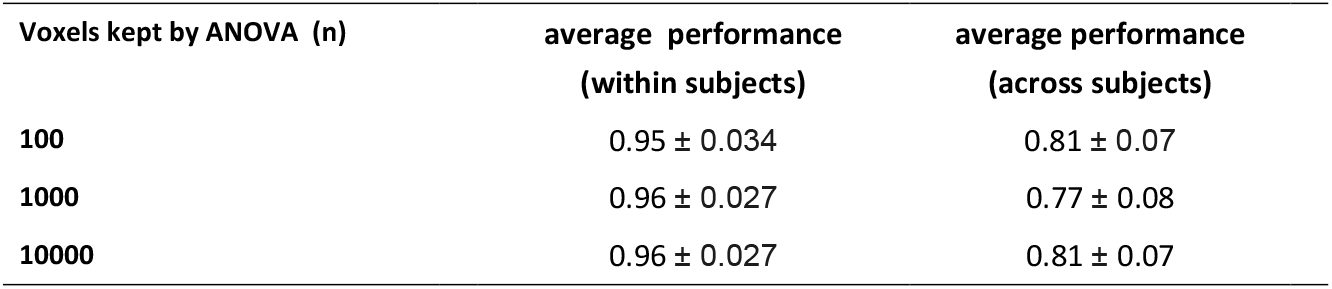
Influence of the size of the mask.

III. Sanity-check analyses:
| Supplementary Table 1. Influence of the size of the mask |  |  |
| --- | --- | --- |
| Voxels kept by ANOVA (n) | average performance<br>(within subjects) | average performance<br>(across subjects) |
| 100 | 0.95 ± 0.034 | 0.81 ± 0.07 |
| 1000 | 0.96 ± 0.027 | 0.77 ± 0.08 |
| 10000 | 0.96 ± 0.027 | 0.81 ± 0.07 |

**Supplementary Figure 2.**
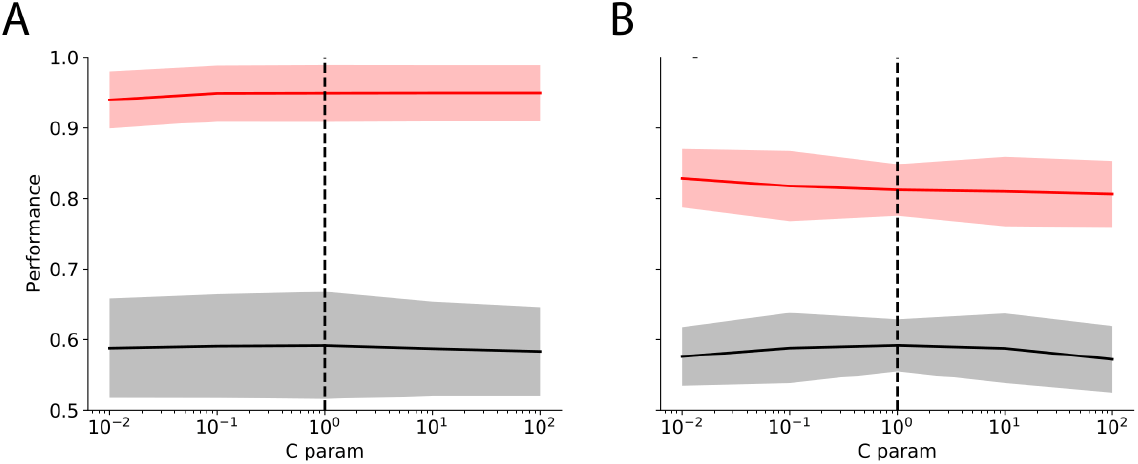
Optimization of the *support vector machine* hyper-parameters. **(A)** Average accuracy over all the subjects enrolled in the **SCZ#1** sample plotted as a function of the C parameter values (see **Methods**). The solid line shows the mean performance, and the shaded areas represent the standard error. The dash-dotted line indicates the value for c=1, chosen in our analyses. **(B)** Same but for the intersubject classification performances.

**Supplementary Figure 3.**
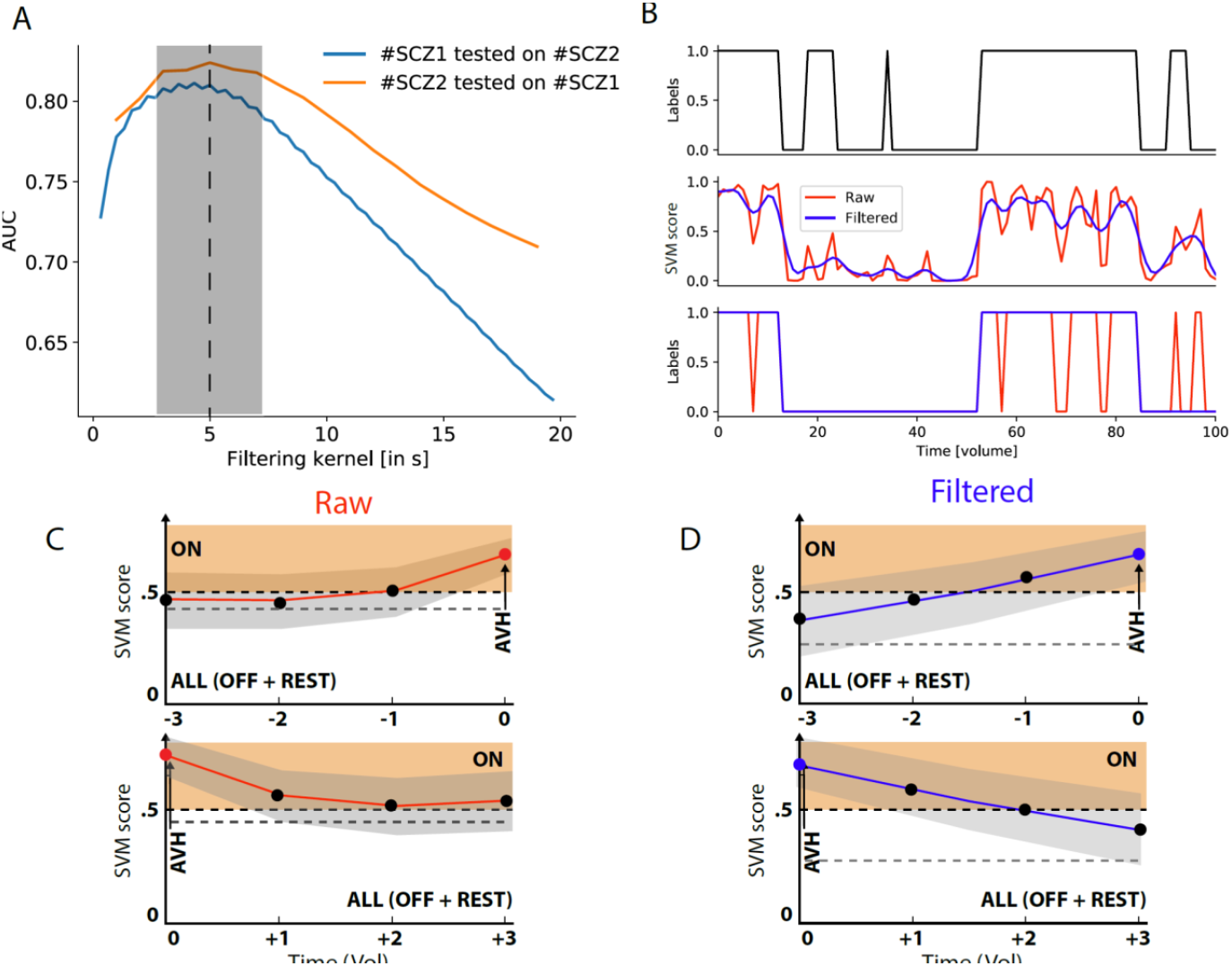
Optimization of the filtering kernel used for real-world application. **(A)** The area under the ROC curve (AUC) is plotted as function of the filtering kernel in seconds to convolve with the raw lSVM scores (see **Methods**) used when the lSVM classifiers are trained on **SCZ#1** (ON vs OFF labels) and tested on all volumes of **SCZ#2** (ON vs ALL) (blue curve) or conversely, when trained on **SCZ#2** (ON vs OFF labels) and tested on all volumes of **SCZ#1** (ON vs ALL labels). Gray shaded area indicates the region where AUC is higher than 0.8, on average, for both curves (**B)** Illustration of the effects of the smoothing kernel on classification. Top row: AVH periods correspond to a y (labels) value of 1. Middle row: SVM score values above 0.5 indicate AVH detection (± SEM). SVM scores either using the raw output (red curve), or the filtered output with a kernel of 5s (blue curve), for **SCZ#1** tested on all volumes of **SCZ#2**, equivalent to a temporal smoothing. Bottom row: final decision either with (blue curve) or without filtering (red curve). **(C - D) Cross decoding AVH before (upper panel) and after their onset (lower panel).** To get a better sense of the temporal specificity of the classifiers, we applied the decoder trained with ON/OFF labels to time points preceding and following AVH. We can see that for the volumes close to the AVH (either before or after), the high score of the SVM can potentially lead to False Positives. decoder based on raw data is plotted in red **(C)**, while the filtered one is shown in blue **(D)**.

**Supplementary Figure 4.**
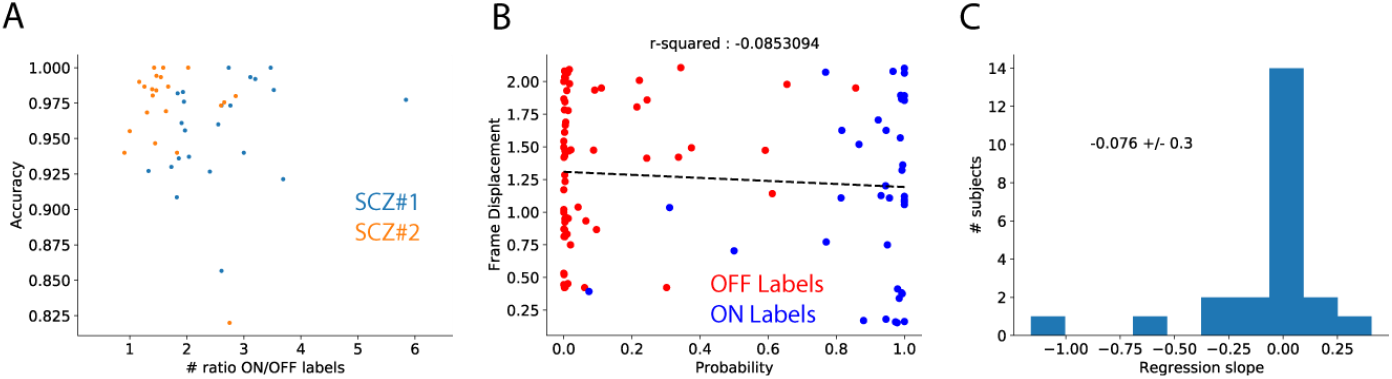
Complementary analyses of the classfier’s robustness. **(A)** Absence of association between hallucinatory occurrences in a given patient (nb of ON labels, 1 pt is 1 subject) and the classifier’s performance for the two datasets (**SCZ#1** and **SCZ#2**). (B) Absence of association between motion artifacts in a given subject (frame displacements (FD), see **Methods**) and the classification performances for ON/OFF label detection (e.g., subject #24). The dash-dotted line indicates the linear regression between the two quantities (r^2^=-0.02, p=0.6). (C) Distribution of the regression slope values between FD and the classification performance (as shown in (B), computed for all the subjects enrolled in the **SCZ#1** sample, confirming an absence of association (mean value ~ 0.076).

### IV. AVH-decoder’s performances using a noisier dataset

**Supplementary Figure 5.**
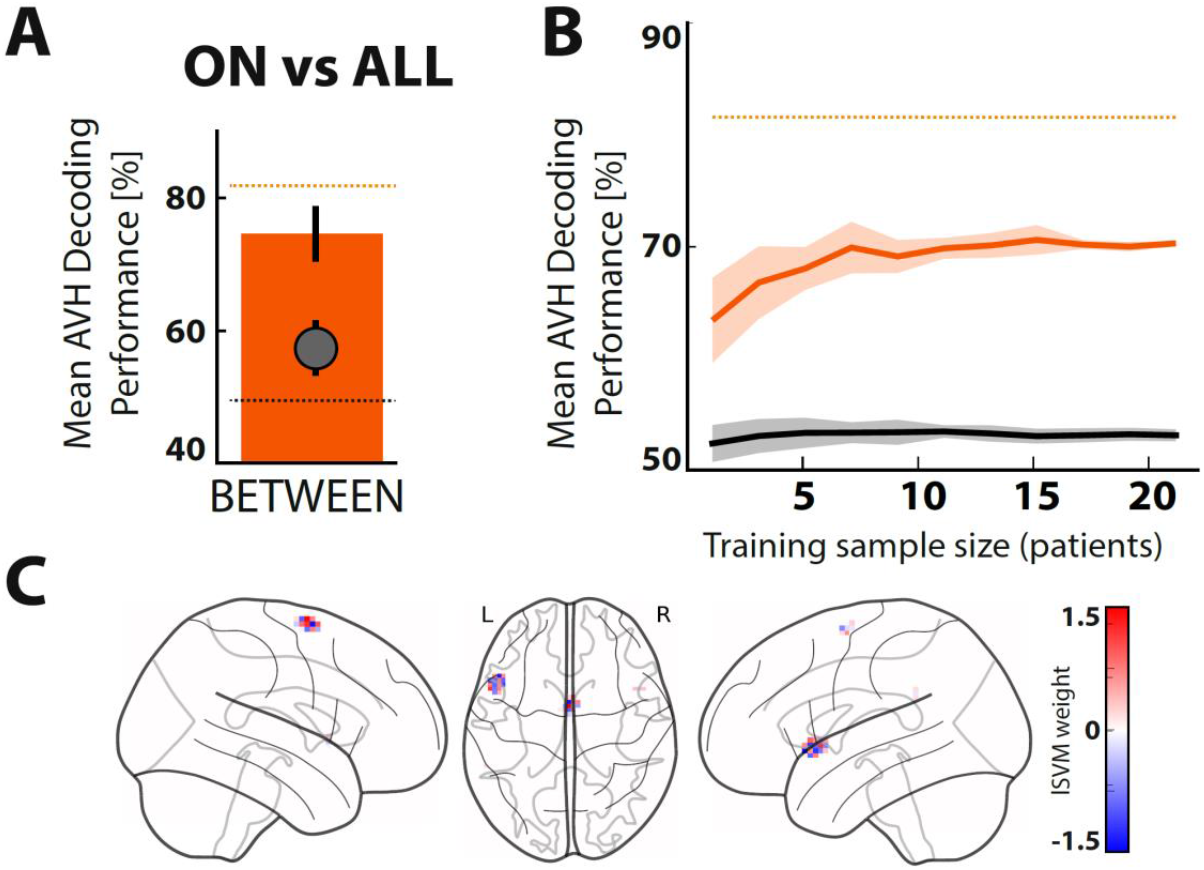
Reliability and performances of decoding auditory-verbal hallucinations (AVH) between-subjects using a lSVM classifier (ON *vs*. ALL=OFF+REST). **(A) Group average of between-subject AVH decoding performance** (ON vs. ALL, **SCZ#1** dataset). The black circle indicates chance level, as estimated using Monte Carlo simulation (with 1000 permutations). Error bars indicate between-subject standard-error to the mean (SEM). **(B) Reliability of between-subjects AVH decoding performance.** Mean between-subjects decoding performance of ON periods against OFF and REST periods (ALL) as a function of the number of patients in the training-set (red line, **SCZ#1** dataset). The black line indicates chance performance level estimated using Monte Carlo simulation (1000 permutations). Error bars indicate SEM. **(C) Contribution map**. The one hundred most informative voxels are color coded to illustrate their contribution to the classifier (lSVM weight). L: left; R: right side of the brain.

### V. ALE meta-analysis of per-hallucinatory functional imaging studies

**Supplementary Table 2.** Characteristics of the studies included in the meta-analysis measuring functional brain activity associated with auditory verbal hallucinations (AVH) in schizophrenia.

| Author | Imaging modality | Pop | Diag criteria | Design | Nb of foci | Original stereotaxic space |
| --- | --- | --- | --- | --- | --- | --- |
| Copolov et al, 2003 | PET | 8 SCZ or SCA | DSM-IV | Online button-press during AVH | 6 | TAL |
| Diederen et al, 2013 | fMRI | 63 SCZ | DSM-IV | Online button-press during AVH | 5 | MNI |
| Diederen et al, 2012 | fMRI | 21 SCZ | DSM-IV | Online button-press during AVH | 7 | MNI |
| Dierks et al, 1999 | fMRI | 3 SCZ | DSM-III-R | Online button-press during AVH | 22 | TAL |
| Horga et al, 2014 | PET | 9 SCZ | DSM-IV | Random-sampling method | 1 | MNI |
| Jardri et al, 2007 | fMRI | 1 COS | DSM-IV-R | ICA + a-posteriori interview | 4 | TAL |
| Jardri et al, 2009 | fMRI | 1 SCZ | DSM-IV-R | ICA + online button-press during AVH | 10 | TAL |
| Jardri et al, 2013 | fMRI | 20 FEP | DSM-IV-R | ICA + a-posteriori interview | 11 | TAL |
| Lefebvre et al, 2016 | fMRI | 20 SCZ | DSM-IV-R | ICA + a-posteriori interview | 16 | MNI |
| Lennox et al, 2000 | fMRI | 4 SCZ | ICD-10 | Online button-press during AVH | 19 | TAL |
| Leroy et al, 2017 | fMRI | 20 SCZ | DSM-IV-R | ICA + a-posteriori interview | 19 | TAL |
| Mallikarjun et al, 2018 | fMRI | 18 FEP | DSM-IV | Online button-press during AVH | 15 | MNI |
| Raij et al, 2009 | fMRI | 11 SCZ | DSM-IV | Online button-press during AVH | 11 | TAL |
| Shergill et al, 2000 | fMRI | 6 SCZ | DSM-IV | Random-sampling method | 20 | TAL |
| Shergill et al,<br>2001 | fMRI | 1 SCZ | DSM-IV | Random-sampling<br>method | 3 | TAL |
| Silbersweig et<br>al, 1995 | PET | 5 SCZ | DSM-IV | Online button-press<br>during AVH | 9 | TAL |
| Sommer et al,<br>2008 | fMRI | 24 SCZ<br>or SCA | DSM-IV | Online button-press<br>during AVH | 21 | MNI |
| Thoma et al,<br>2016 | fMRI | 16 SCZ<br>or SCA | DSM-IV-R | Online button-press<br>during AVH | 9 | MNI |

fMRI: functional magnetic resonance imaging; PET: positron emission tomography; SCZ: schizophrenia; COS: childhood-onset schizophrenia; FEP: first-episode of psychosis; SCA: schizo-affective disorder; DSM: diagnostic statistical manual for psychiatric disorders; ICD: WHO international classification of diseases; TAL: Talairach & Tournoux standardized space; MNI: Montreal Neurological Institute standardized space; ICA: independent component analysis.

**Supplementary Table 3.** Brain regions with significantly elevated likelihoods of activation during auditory-verbal hallucinations.

| Gyri | Contribution | MNI coord (x/y/z) |  |  | ALE | Z | P |
| --- | --- | --- | --- | --- | --- | --- | --- |
| Superior temporal gyrus (STG) | 23% | -52 | 10 | -2 | 0.0193 | 5.0133 | 2.6752 e-7 |
| Anterior Insula + Inferior frontal gyrus | 52% (AI) + 14% (IFG) | -44 | 4 | -4 | 0.0152 | 4.2735 | 9.6189 e-6 |
| Precentral gyrus | 11% | -56 | 4 | 10 | 0.0144 | 4.1284 | 1.8260 e-5 |
| Anterior insula + STG | 52% (AI) + 26% (STG) | 52 | 12 | -4 | 0.0175 | 4.7124 | 1.2241 e-6 |
| // | // | 48 | 8 | -10 | 0.0141 | 4.0454 | 2.6117 e-5 |
| Inferior Frontal Gyrus + Precentral gyrus | 18% (IFG) + 3% (PCG) | 56 | 20 | -10 | 0.0134 | 3.9256 | 4.3257 e-5 |

The inferior frontal gyrus gathered Brodmann's area 44 and 47.

**Supplementary Figure 6.**
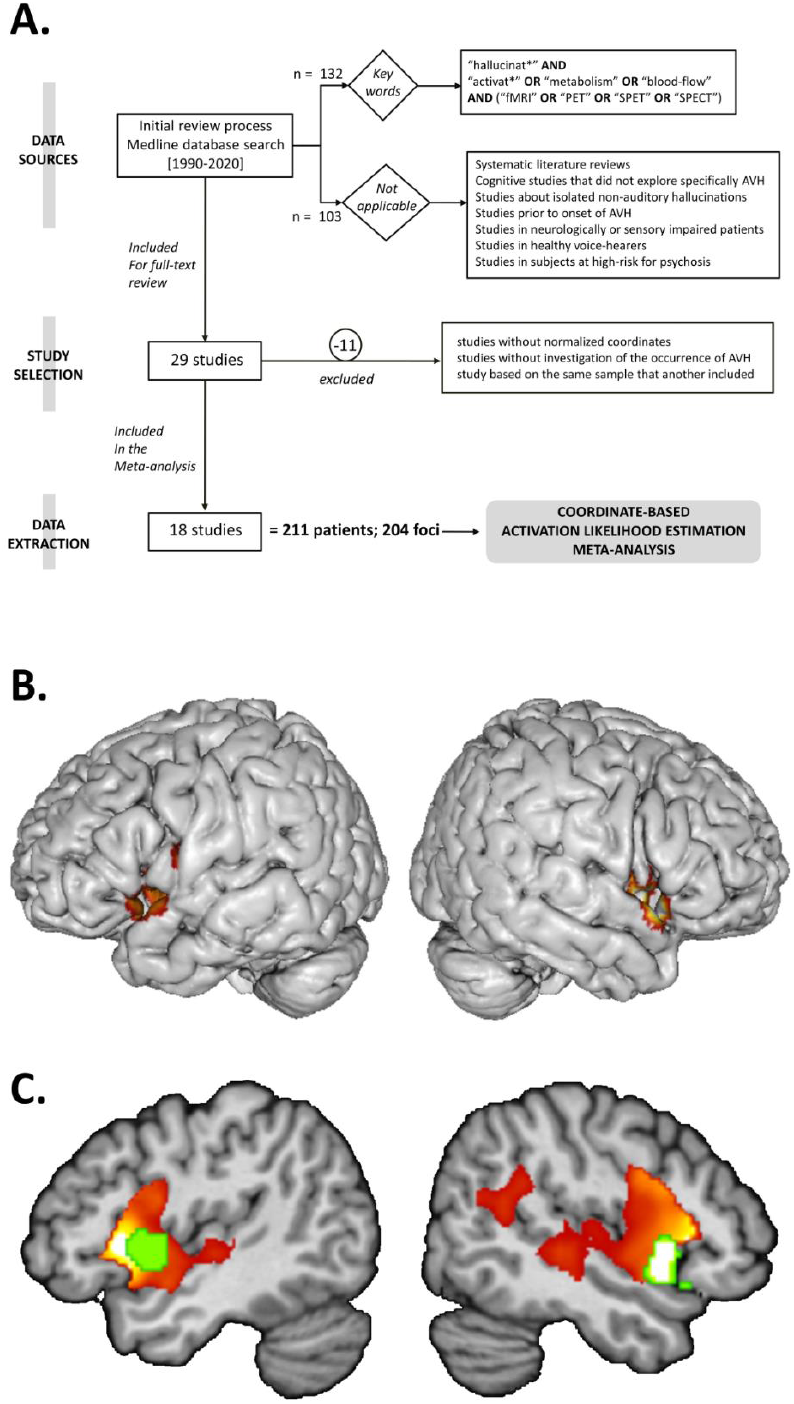
Coordinate-based meta-analysis (CBMA) of per-hallucinatory functional brain imaging studies. We present an updated 2020 Activation Likelihood Estimation (ALE) map of the seminal Jardri et al. meta-analysis (11). We referred to an identical search algorithm on the Medline database and identified 132 papers for the period 1990 - 2020. After applying selection criteria summarized in the PRISMA flow-chart presented in **(A)**, 18 studies were retained for meta-analysis (see also **Suppl. Table 3**), gathering 211 hallucinators with schizophrenia and 204 per-AVH foci. Using GingerALE v 3.02 software, we ran a random-effect CBMA. Following current recommendations, we used a cluster-level FWE threshold of 0.05 with 1000 permutations (the cluster-forming threshold at the voxel level was fixed at p< 0.001). **(B)** Two clusters centered on the right/left Broca areas were identified during AVH occurrences (Cf. also **Suppl. Table 3**, map available at NeuroVault #). These clusters (in green in panel C) overlap with the contributive regions identified in the present paper (Cf. **Figure 5B**), as well as with overactivated areas previously identified in individuals prone to hear voices during an induced auditory hallucinations task shown in yellow-to-red in panel **(C)** (32).

